# Safe, effective, and inexpensive clearance of mycoplasma contamination from cultures of apicomplexan parasites with Sparfloxacin

**DOI:** 10.1101/2022.08.30.505931

**Authors:** Darlene R. Malave-Ramos, Kit Kennedy, Melanie N. Key, Zhicheng Dou, Björn F.C. Kafsack

## Abstract

Most commercial products cannot be used for clearance of mycoplasma contamination from cultures of apicomplexan parasites due to the parasites’ dependence on the apicoplast, an essential organelle with DNA replication and translation machinery of cyanobacterial origin. The lone exception, Mycoplasma Removal Agent (MRA), is relatively expensive and some mycoplasma strains have shown resistance to clearance with MRA. Here, we report that the fluoroquinolone antibiotic Sparfloxacin is a safe, effective, and inexpensive alternative for treatment of mycoplasma contamination in cultures of apicomplexan parasites. Sparfloxacin cleared both MRA- sensitive and MRA-resistant mycoplasma species from *P. falciparum* cultures at 1 and 4 μg/mL, respectively. We show that cultures of three different apicomplexan parasites can be maintained at concentrations of Sparfloxacin required to clear mycoplasma without resulting in substantial deleterious effects on parasite growth. We also describe an alternative low-cost, in-house PCR assay for detecting mycoplasma. These findings will be useful to laboratories maintaining apicomplexan parasites in vitro, especially in low-resource environments, where the high cost of commercial products creates an economic barrier for detecting and eliminating mycoplasma from culture.

## INTRODUCTION

Contamination with mycoplasma species (class Mollicutes) is a common problem in continuous cell culture and found in 15-35% of cell lines ^1–3^, including co-cultures of apicomplexan parasites and their host cells. Dozens of mycobacterial species have been found as part of the human microbiota, including skin and hair, but only a small number have been linked to human disease. With a 0.15-0.3 micron diameter, mycoplasmas are some of the smallest known cellular organisms and can pass through the standard 0.2 micron filters standardly used in the preparation of culture media. Contamination of cell cultures with mycoplasma is easily missed for several reasons, including their small size, poor staining due to the absence of a cell wall, and failure to change culture turbidity or pH ^4^. Unlike contamination with most other bacteria or fungi, the presence of mycoplasma rarely results in the collapse of the cell culture but can nonetheless substantially alter the properties and behavior of infected cell lines, making it a potentially important confounder in experimental comparisons.

Mycoplasma species are broadly resistant to many classes of commonly used antibiotics, including beta-lactams, glycopeptides, fosfomycin, polymixins, sulfonamides, rifampicin, first-generation quinolones, and trimethoprim. Conversely, they remain susceptible to some tetracyclines, macrolides, and second-generation quinolones ^5^. These classes form the basis of commercial products aimed at eradicating mycoplasma contamination from cell cultures, such as Bayer’s Ciprobay and Promocell’s Biomyc-3 (both the fluoroquinolone Ciprofloxacin), Promocells’s Biomyc-1/2 and Roche’s BM-Cyclin (both are a combination of the macrolide Tiamulin and the tetracycline Minocycline), Invivogen’s Plasmocin (a combination of a proprietary fluoroquinolone and a proprietary macrolide), Bayer’s Baytril (the fluoroquinolone Enrofloxacin), and MP Biomedicals’ Mycoplasma Removal Agent (a proprietary fluoroquinolone).

Unfortunately, nearly all these products also inhibit the growth of *Plasmodium falciparum* ^6–8^ or other apicomplexan parasites ^9,10^ at similar or lower concentrations than those required to effectively clear mycoplasma ^11–14^. This poor selectivity is due their effect on the parasite apicoplast, an essential chloroplast-derived organelle that retains translation and DNA-replication machinery of cyanobacterial origin. The lone validated exception is Mycoplasma Removal Agent (MRA), which contains an undisclosed fluoroquinolone and has successfully been used to clear mycoplasma from cultures of apicomplexan parasites, including *P. falciparum* ^6^, *Toxoplasma gondii,* and *Neospora caninum* ^15^. Use of MRA has some downsides; clearance of mycoplasma with MRA can take 1-3 weeks at the recommended concentration of 0.5 μg/mL, and some mycoplasma strains have shown resistance to clearance with MRA ^12,13^. Furthermore, MRA is relatively expensive at $270 per liter of culture medium.

Here we report that the fluoroquinolone Sparfloxacin is a safe, effective, and low-cost alternative to MRA for treatment of mycoplasma contamination. Sparfloxacin cleared both MRA-sensitive and MRA-resistant mycoplasma species from *P. falciparum* cultures at 1 and 4 μg/mL, respectively. Furthermore, we show that cultures of three different apicomplexan parasites can be maintained without deleterious effects on parasite growth at concentrations of Sparfloxacin required to clear mycoplasma.

## RESULTS & DISCUSSION

### Dose-response of Apicomplexan parasites to Sparfloxacin

To establish the maximal concentrations of Sparfloxacin that cultures of apicomplexan parasites would tolerate, we measured the effect of Sparfloxacin on the growth of three geographically diverse strains of *Plasmodium falciparum* (Figure 1a), as well as *Toxoplasma gondii* and *Babesia divergens (*Figure 1b), two other commonly cultured apicomplexan parasites. EC50 values of the tested isolates all fell within a narrow range of 3.1-8.6 μg/mL.

**Figure 1.**
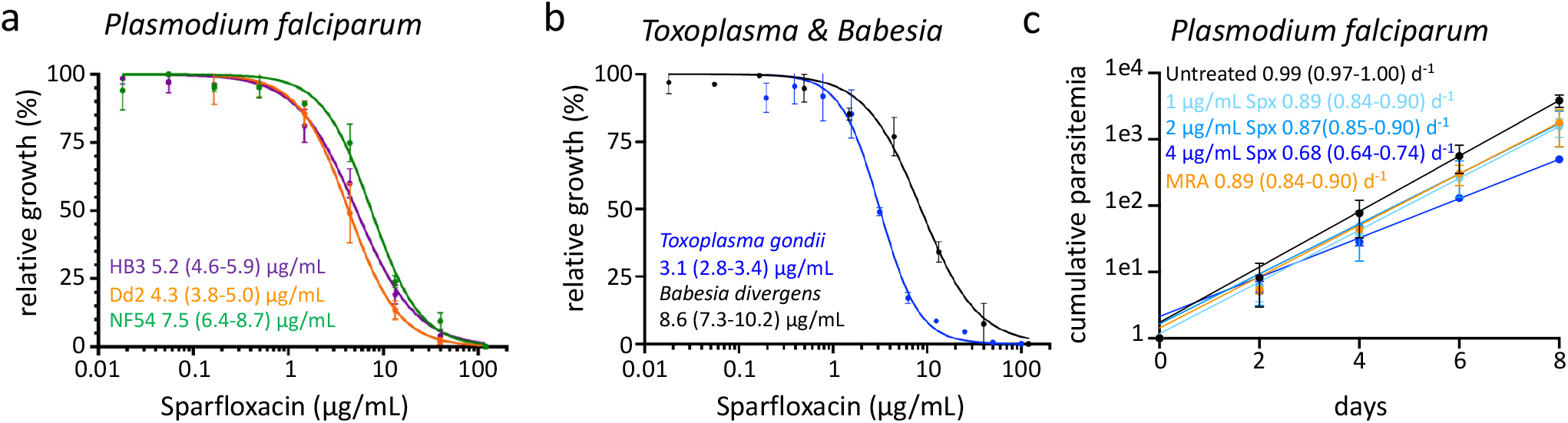
Dose-response of apicomplexan parasites to Sparfloxacin. **a)** Sparfloxacin dose-response of *Plasmodium falciparum* isolates of African (NF54, green), Southeast Asian (Dd2, orange) and South American (HB3, blue) origin based on 72-hour SYBR Green growth assays. Estimated EC50 with 95% confidence are indicated based on n=2-3. **b)** Sparfloxacin dose-responses of *B. divergens* isolate Rouen 1987 and *T. gondii* were measured using on 72-hour SYBR Green or 96-hour luciferase growth assays, respectively. Estimated EC50s with 95% confidence are indicated based on n=2-3. **c)** Representative growth curves of *P. falciparum* NF54 maintained at 1/2/4 μg/mL Sparfloxacin, MRA (0.5 μg/mL), or no drug. Growth rates with 95% confidence interval are shown in days^-1^ based on n=2.

The effect of drugs inhibiting apicoplast replication doesn’t manifest until parasites have invaded their next host cell, a phenomenon known as “delayed death” ^16^. Since the growth assays used for *Toxoplasma* and *Babesia* parasites covered multiple lytic cycles (Figure 1b), these results already capture any delayed death effects of Sparfloxacin on apicoplast replication. However, since *P. falciparum* divides by schizogony and apicoplast replication only occurs every 48 hours, our 72 hour growth assays may not have fully captured any potential delayed death effects by Sparfloxacin. To evaluate whether *P. falciparum* cultures exhibited any delayed death, we measured the effect of Sparfloxacin over 4 replication cycles (Figure 1c). Growth rates of *P. falciparum* NF54 cultures were constant over 8 days at all concentrations of Sparfloxacin, excluding the possibility of substantial delayed death effects, which would have resulted in a decrease of the growth rate after the first replication cycle. Compared to untreated cultures, there was a 10-12% decrease in the growth rate when parasites were maintained in the presence of 1 and 2 μg/mL of Sparfloxacin or 0.5 μg/mL MRA. Growth slowed by 31% at 4 μg/mL but cultures could be maintained continuously for at least 3 weeks at this concentration (Figure 3b). While we did not quantify the effect on continuous growth, all the other *P. falciparum* strains, as well as *Toxoplasma* and *Babesia,* could be maintained at 4 μg/mL for prolonged periods.

### A low-cost, in-house PCR assay with internal controls is effective at detecting mycoplasma contamination

We compared the ability to detect mycoplasma in DNA isolated from cultures of *P. falciparum* for an in-house PCR assay, based on universal mycoplasma primers targeting 425 bp of the locus encoding the 16S ribosomal RNA ^17^, to a commercially available mycoplasma detection PCR kit. To provide an internal PCR control for negative mycoplasmas results, we included small amount of pUC18 plasmid and primers to amplify 316 bp of the plasmid backbone in the PCR master mix. The commercial PCR product was able to detect mycoplasma in DNA extracted from mycoplasma-positive cultures down to a 1:1000 dilution at a per-sample cost of $6.15. The in-house PCR assay was also effective at detecting mycoplasma at a per-sample cost of $0.20 but the limit of detection was slightly lower at between 1:100 and 1:1000 (Figure 2).

**Figure 2.**
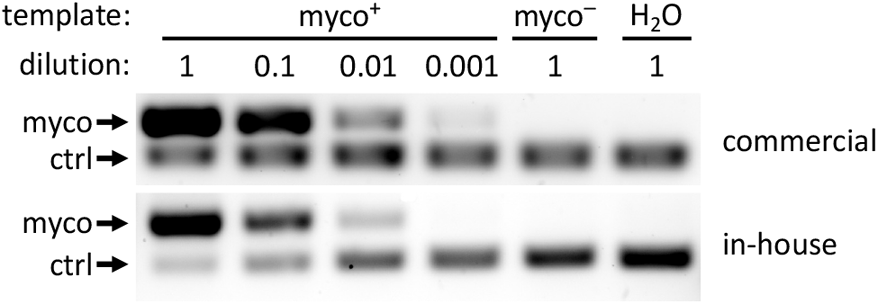
Detection of mycoplasma in *P. falciparum* cultures with commercial or in-house PCR assays. A commercial PCR assay was able detect mycoplasma in DNA of mycoplasma-positive (myco^+^) cultures even at 1:1000 dilution while the limit of detection for the in-house assay was between 1:100 and 1:1000. Results from undiluted mycoplasma-negative (myco^-^) or no template (H2O) are shown as controls. Images are representative of n=2.

**Figure 3.**
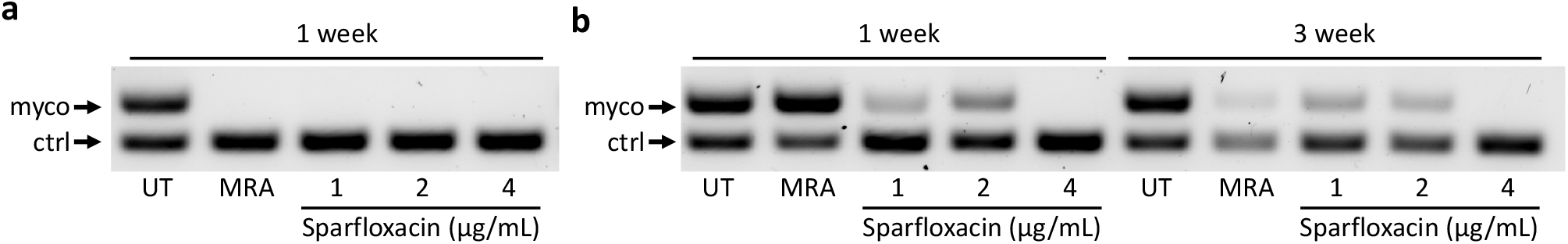
Effective clearance of mycoplasma with Sparfloxaxin. **a)** Sparfloxacin was able to reduce MRA-sensitive mycoplasmas below the level of detection of the in-house PCR assay after one week of treatment at all concentrations tested. **b)** Sparfloxacin was able to clear MRA-resistant mycoplasma species from *P. falciparum* cultures after one week of treatment at 4 μg/mL when MRA at the recommended 0.5 μg/mL failed even after 3 weeks of treatment. Mycoplasma was detected using the in-house PCR assay and images are representative of n=2.

### Sparfloxacin is effective at clearing both MRA-sensitive and MRA-resistant mycoplasma strains

To evaluate the ability of Sparfloxacin to clear mycoplasma, mycoplasma-positive cultures were maintained either without treatment, with the manufacturer recommended 0.5 μg/mL of MRA, or with 1, 2, or 4 μg/mL of Sparfloxacin. For most cultures, one week of treatment with MRA and all concentrations of Sparfloxacin was able to reduce mycoplasma below the level of detection of the in-house PCR assay (Figure 3a). However, mycoplasma contamination in one culture proved to be resistant to clearance with MRA even after three weeks of treatment but was successfully cleared after one week of treatment with 4 μg/mL Sparfloxacin (Figure 3b).

This demonstrates that Sparfloxacin can be used to treat even MRA-resistant mycoplasma strains at concentrations that are well tolerated by a variety of apicomplexan parasites at a cost of $0.06 per liter of culture medium, making it a safe, effective, and low-cost alternative to MRA with its cost of $696 per liter of culture medium.

## MATERIALS & METHODS

### Parasite strains and maintenance

*Plasmodium falciparum* strains NF54, Dd2, and HB3 and *Babesia divergens* strain Rouen 1987 ^18^ were maintained using standard malaria culturing techniques ^19^. *Toxoplasma gondii* strain RHΔku80::NLuc were maintained in Human Foreskin Fibroblasts (ATCC SCRC-1041) using standard culturing techniques ^20^. Briefly, HFF/*T. gondii* co-cultures were maintained at 37°C with 5% CO_2_ in D10 medium (Dulbecco’s Modified Eagle Medium, 4.5 g/L glucose, VWR) supplemented with 10% Cosmic Calf serum (HycloneTM, GE Healthcare Life Sciences SH30087.03), 10 mM HEPES, additional 2 mM L-glutamine, and 10 mM Pen/Strep.

### Mycoplasma detection

Mycoplasma contamination was detected using the e-Myco Mycoplasma detection kit (LiliF Diagnostics) according to manufacturer instructions. Alternatively, mycoplasma was detected using a custom PCR using Taq polymerase manufacturer instructions and 0.4 μM degenerate universal mycoplasma detection primers (GTGGGGAGCAAAYAGGATTAGA/GGCATGATGATTTGACGTCRT) that target the 16S locus ^17^. As an internal control, each reaction also contained 100 fg of pUC18 plasmid and 0.08 μM of primers (CCTGACGAGCATCACAAAAA / AGTCGTGTCTTACCGGGTTG) that amplify a 316 bp section of the plasmid backbone. PCR master mix reagents (see Figure 4): *Taq* polymerase and 10X Reaction Buffer (NEB M0273S), nucleotide mix containing 10 mM dTTP, dATP, dGTP, dCTP (NEB N0446S) each, and 80 fg of pUC18 plasmid. pUC 18 can be substituted with any other pBlueScript II-derived plasmid. 20 μL aliquots of master mix were stored at −20 °C until use.

**Figure 4:**
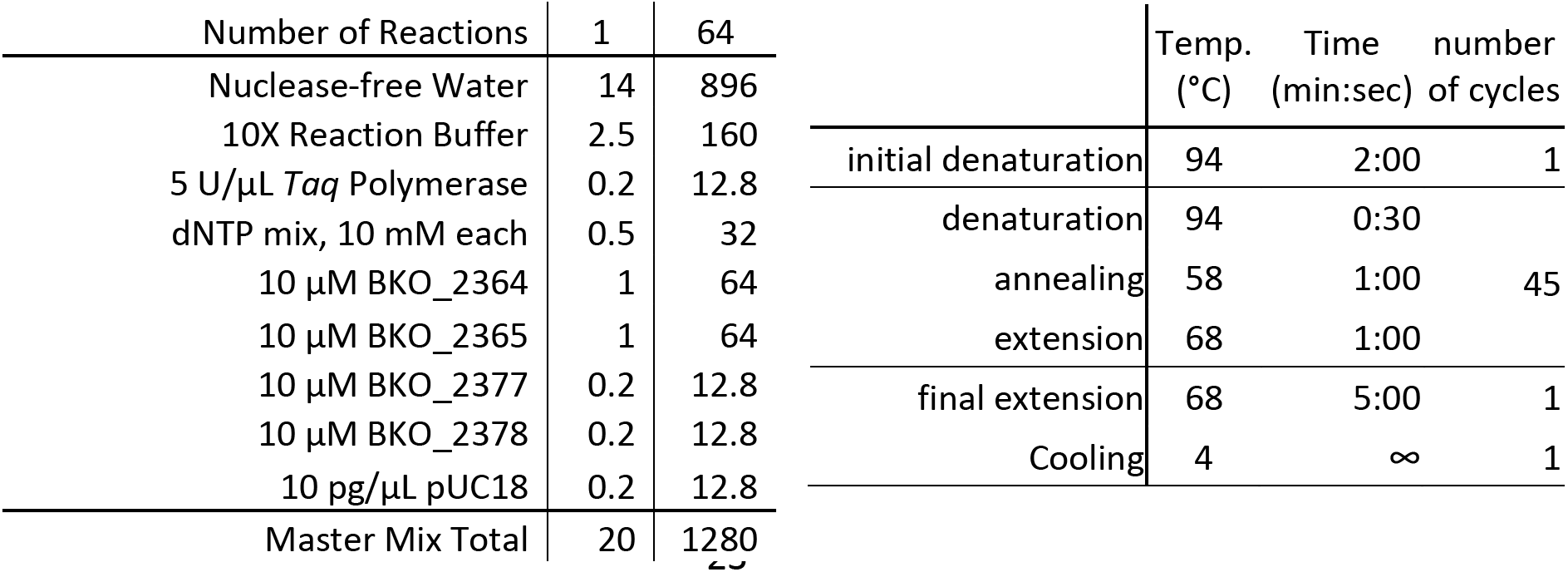
Mycoplasma Detection PCR master mix composition and cycling conditions.

### Dose-response assays

Sparfloxacin (ThermoScientific J66358-09) was dissolved in DMSO to a stock concentration of 2 mg/mL. For *P. falciparum* and *B. divergens* asexual blood-stages the dose response to Sparfloxacin was determined using the 72-hour SYBR Green Growth Assay ^21^ as described in detail in ^22^. For *T. gondii,* 1,500 *RHΔku80::NLuc* tachyzoites were inoculated into each well of a 96-well plate containing confluent HFF cells and the dose response was determined by a 96-hour luciferase-based growth assay as described in detail in ^20^. In brief, the NanoLuc luciferase-expressing *Toxoplasma* parasites were allowed to invade in human foreskin fibroblasts for 4 hrs. Non-invaded parasites were washed off and the media were replaced with replaced with fresh D10 supplemented with Sparfloxacin. Ten concentrations generated by 2-fold serial dilution were tested starting at 100 μg/mL. After 96-hour incubation, the media were gently aspirated and the infected host cells were treated in the lysis buffer (100 mM 4- Morpholineethanesulfonic acid (MES) pH 6.0, 1 mM trans-1,2-Diaminocyclohexane-N,N,N’,N’-tetraacetic acid (CDTA), 0.5% (v/v) Tergitol, 0.05% (v/v) Mazu DF 204, 150 mM KCl, 1 mM DTT, and 35 mM thiourea) containing 12.5 μM Coelenterazine H and incubated at room temperature for 10 min prior to luminescence quantification using a BioTek Synergy H1 Hybrid plate reader. The luminescence signals from Sparfloxacin-treated cells were normalized against that from infected cells grown in plain D10 medium. The IC50 concentration for each species was estimated by non-linear fit using Prism (GraphPad).

### Effect of Sparfloxacin on *P. falciparum* growth rate

Growth of *P. falciparum* cultures were followed by measuring culture parasitemia every other day for 8 days by flow cytometry after staining live cultures with 16 μM Hoechst 33342 and 200 nM Thiazole Orange for 30 min at 37°C. Cultures above 2% were diluted to 0.5% and actual dilution factor was determined by measuring parasitemia again after dilution using flow cytometry. Cumulative parasitemia was calculated by multiplying the percentage of infected cells by the overall dilution factor and the growth rate was determined by exponential fit using Prism (GraphPad).

## Author contributions

Conceptualization: BFCK, KK; Methodology: BK, KK, ZD; Investigation: DRMR; Writing – Original Draft: BFCK Writing – Review & Editing: BFCK, KK; Visualization: KK, BFCK; Supervision: BFCK, KK, ZD; Project Administration: BFCK; Funding Acquisition: BFCK, ZD.

## Data and materials availability

No new materials were generated in this study.

## REFERENCES

1. McGarrity, G. J. & Barile, M. F. Methods in Mycoplasmology V2: Diagnostic Mycoplasmology. (1983).

2. Drexler, H. G. & Uphoff, C. C. Mycoplasma contamination of cell cultures: Incidence, sources, effects, detection, elimination, prevention. in 39, 75–90 (2002).

3. McGarrity, G. J., Phillips, D. M. & Vaidya, A. B. Mycoplasmal infection of lymphocyte cell cultures: Infection with M. salivarium. In Vitro 16, 346–356 (1980).

4. Rottem, S. Interaction of mycoplasmas with host cells. Physiological Reviews 83, 417–432 (2003).

5. Gautier-Bouchardon, A. V. Antimicrobial Resistance in Mycoplasma spp. Microbiol Spectr 6, (2018).

6. Singh, S., Puri, S. K. & Srivastava, K. Treatment and control of mycoplasma contamination in Plasmodium falciparum culture. Parasitol Res 104, 181–184 (2008).

7. Mahmoudi, N. et al. In vitro activities of 25 quinolones and fluoroquinolones against liver and blood stage Plasmodium spp. Antimicrobial Agents and Chemotherapy 47, 2636–2639 (2003).

8. Pradines, B. et al. In vitro activities of antibiotics against Plasmodium falciparum are inhibited by iron. Antimicrobial Agents and Chemotherapy 45, 1746–1750 (2001).

9. Rizk, M. A. et al. Inhibitory effects of fluoroquinolone antibiotics on Babesia divergens and Babesia microti, blood parasites of veterinary and zoonotic importance. Infect Drug Resist 11, 1605–1615 (2018).

10. Batiha, G. E.-S. et al. Inhibitory effects of novel ciprofloxacin derivatives on the growth of four Babesia species and Theileria equi. Parasitol Res 119, 3061–3073 (2020).

11. Bébéar, C. M., Renaudin, H., Schaeverbeke, T., Leblanc, F. & Bébéar, C. In-vitro activity of grepafloxacin, a new fluoroquinolone, against mycoplasmas. J Antimicrob Chemother 43, 711–714 (1999).

12. Drexler, H. G. et al. Treatment of mycoplasma contamination in a large panel of cell cultures. In Vitro Cell Dev Biol Anim 30A, 344–347 (1994).

13. Uphoff, C. C., Denkmann, S.-A. & Drexler, H. G. Treatment of mycoplasma contamination in cell cultures with Plasmocin. Journal of Biomedicine and Biotechnology 2012, 267678 (2012).

14. Molla Kazemiha, V. et al. Effectiveness of Plasmocure™ in Elimination of Mycoplasma Species from Contaminated Cell Cultures: A Comparative Study versus other Antibiotics. Cell J 21, 143–149 (2019).

15. Hudson, A. & Ellis, J. T. Culture of Neospora caninum in the presence of a Mycoplasma Removal Agent results in the selection of a mutant population of tachyzoites. Parasitology 130, 607–610 (2005).

16. Kennedy, K., Crisafulli, E. M. & Ralph, S. A. Delayed Death by Plastid Inhibition in Apicomplexan Parasites. Trends in Parasitology 35, 747–759 (2019).

17. Molla Kazemiha, V. et al. PCR-based detection and eradication of mycoplasmal infections from various mammalian cell lines: A local experience. Cytotechnology 61, 117–124 (2009).

18. Gorenflot, A. et al. Cytological and immunological responses to Babesia divergens in different hosts: ox, gerbil, man. Parasitol Res 1–10 (1991).

19. Schlichtherle, M. Malaria Research and Reference Reagent Resource Center. Methods in Malaria Research. (2000).

20. Key, M. et al. Determination of Chemical Inhibitor Efficiency against Intracellular Toxoplasma Gondii Growth Using a Luciferase-Based Growth Assay. JoVE (Journal of Visualized Experiments) 2020, (2020).

21. Smilkstein, M., Sriwilaijaroen, N., Kelly, J. X., Wilairat, P. & Riscoe, M. Simple and inexpensive fluorescence-based technique for high-throughput antimalarial drug screening. Antimicrobial Agents and Chemotherapy 48, 1803–1806 (2004).

22. Kirkman, L. A. et al. Antimalarial proteasome inhibitor reveals collateral sensitivity from intersubunit interactions and fitness cost of resistance. Proceedings of the National Academy of Sciences 115, E6863–E6870 (2018).

